# Automated 3D Neuron Tracing with Precise Branch Erasing and Confidence Controlled Back-Tracking

**DOI:** 10.1101/109892

**Authors:** Siqi Liu, Donghao Zhang, Yang Song, Hanchuan Peng, Weidong Cai

**Affiliations:** School of Information Technologies, University of Sydney, Darlington, NSW Australia.; Allen Institute for Brain Science, Seattle, WA, USA.

**Keywords:** 3D neuron reconstruction, Neuron morphology

## Abstract

The automatic reconstruction of single neuron cells from microscopic images is essential to enabling large-scale data-driven investigations in neuron morphology research. However, few previous methods were able to generate satisfactory results automatically from 3D microscopic images without human intervention. In this study, we developed a new algorithm for automatic 3D neuron reconstruction. The main idea of the proposed algorithm is to iteratively track backwards from the potential neuronal termini to the soma centre. An online confidence score is computed to decide if a tracing iteration should be stopped and discarded from the final reconstruction. The performance improvements comparing to the previous methods are mainly introduced by a more accurate estimation of the traced area and the confidence controlled back-tracking algorithm. The proposed algorithm supports large-scale batch-processing by requiring only one hyper-parameter for background segmentation. We bench-tested the proposed algorithm on the images obtained from both the DIADEM challenge and the BigNeuron challenge. Our proposed algorithm achieved the state-of-the-art results.

## I. INTRODUCTION

DIGITAL reconstruction of 3D neuron morphological models is important for understanding the connectivity of the nervous system and the cell information processing within neurons. Given a 3D light microscopic image stack containing a single neuron, the reconstruction resembles a tree graph model that represents the circuit of the neuron cell. Within the scope of computational neuroscience, the reconstructed models are acquired for purposes such as neuronal identity, anatomically and biophysically realistic simulations, morphometric and stereological analysis and determining potential connectivity. On the other hand, the speed of image acquisition techniques has greatly surpassed the speed of processing these images. The 3D neuron models being used for neuron morphology studies nowadays were mainly generated by manual or semiautomatic tracing methods, which is a highly time-consuming task. The automatic reconstruction of the neuron morphological models has thus become one of the core bottlenecks in neuroscience nowadays. The DIADEM challenge [1] and the recent BigNeuron challenge [2] were also hosted to provide open-access data and software tools for improving the accuracy of neuron reconstruction algorithms. However, most automatic tracing methods still tend to fail with low-quality images.

The challenges of neuron reconstruction are mainly caused by the low image quality and the complex neuronal morphology. Due to the fundamental limits of light microscopic imaging and neuron cell extraction pipelines, the 3D microscopic image-stacks often contain strong background noise, irrelevant structures and small gaps along the neuronal arbours. Image qualities from different sites also vastly differ due to the different imaging pipelines.

Many recent methods were proposed to automate 3D neuron reconstruction by combining computer vision techniques and neuron morphological knowledge [3]–[14]. The state-of-theart algorithms are often pipelines combining preprocessing, branch tracing, and post-processing components. According to a recent review paper [15], the existing neuron tracing methods can be divided into global processing [3], [16]–[20], local processing [21]–[23], and meta-algorithms [5], [24], [25]. The global approaches process the entire image whereas the local processing methods explore the image only around the fibres of interests. Some of the meta-algorithms were proposed to tackle the challenges of low image quality or large image scale independently of any specific neuron tracing algorithm. Global processing algorithms are becoming more popular in the recent years than the local processing algorithms since the global information is essential to generate the correct neuronal topology.

The Rivulet algorithm was proposed [8], [26] as a combination of global and local approaches. The global information is firstly explored with the Multi-Stencils Fast Marching [27]. Rivulet then iteratively tracks neuronal arbours from the furthest potential termini back to the soma centre and erases the areas covered by newly traced branches. The branch erasing ensures the algorithm does not generate duplicated arbours. The tracing finishes only when a high proportion of the foreground area has been explored. However, the Rivulet algorithm tends to generate many over-reconstructed arbours and connection errors when the image contains strong noise. The performance of Rivulet is also highly dependent on the choice of three hyper-parameters, which makes it hard to be applied to large-scale datasets.

In this study, we present an algorithm, named Rivulet2, that generates more accurate neuron tracing results with fewer hyper-parameters and faster speed than Rivulet. We refer the original Rivulet algorithm as Rivulet1 for clarity. The major components of the proposed algorithm can be summarised as (1) Preprocessing the image to obtain a segmentation as shown in Fig. 1(b) and generating a distance transform shown in Fig. 1(c); (2) Applying the multi-stencils fast-marching (MSFM) as shown in Fig. 1(d) on the distance transform and computing the gradients of the MSFM time crossing map; (3) Iteratively tracking back from the geodesic furthest point on the foreground and erasing the area that covered by the newly traced branch shown in Fig. 1(e) and (4) Post-processing the result neuron by pruning the short leaves and the unconnected branches to obtain the final results as shown in Fig. 1(f).

**Fig. 1.**
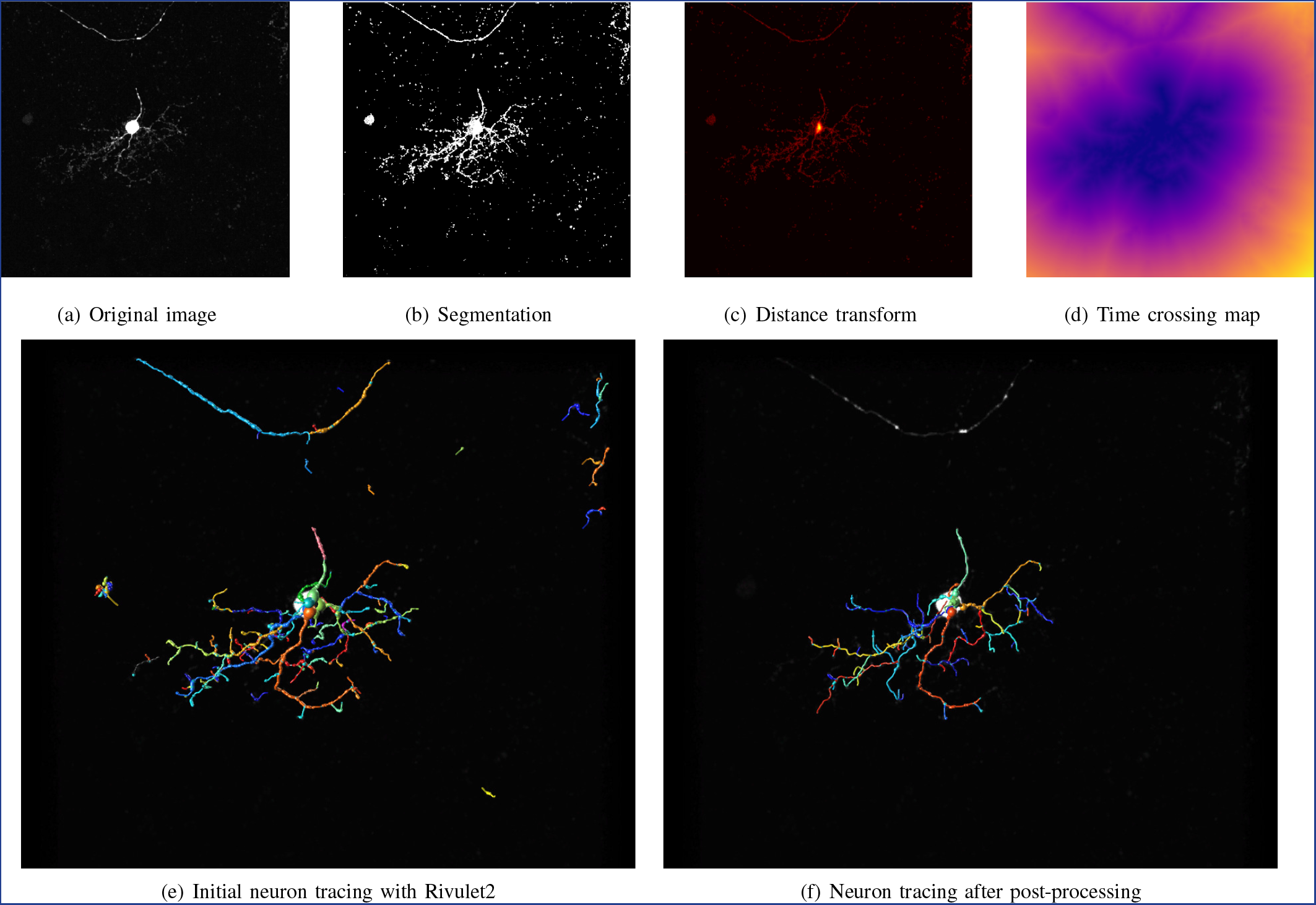
The original image of a zebrafish neuron is shown in (a). Along with the neuron cell of interest, the image also contains many noise and some irrelevant fibres. The example effects of the preprocessing components are shown in (b)-(d). The initial tracing is shown in (e) preserves irrelevant fibres that might be wrongly included in the neuron extraction. The final tracing shown in (f) is obtained by eliminating the redundant fibres and fuzzy leaves. The branch colours are randomised for visualisation.

Comparing to Rivulet1, the novelty of Rivulet2 mainly resides in the (3) and (4) components. The back-tracking of Rivulet1 stops after it traces on the background for a long distance, which is determined by a gap parameter. However, it is ill-posed to set a single hyper-parameter to distinguish the gap distances between unconnected neuronal segments since there could be gaps between noise points and neuronal segments as well. The gap parameter is no longer needed in the the proposed Rivulet2 algorithm since the branch backtracking is stopped by two new hyper-parameter free criteria. The first criterion is computed with an online confidence score that is updated at every tracing step. The second criterion is to check if a large gap presents on an arbour by comparing the gap distance and a score determined by the mean radius sampled at the previous tracing steps. Combing both criteria, Rivulet2 can trace the neuronal arbours with high accuracy even when neuronal segment gaps and strong noise both reside in the image. Rivulet1 also generates small fuzzy branches which are caused by the coarsely estimated neuron surface. We present a method to erase the traced branches precisely for suppressing the false positive branches. It also makes the proposed algorithm faster than the original Rivulet since it finishes within fewer iterations without revisiting the traced image areas. A parameter-free approach is added to merge the newly traced branches into the neuron tree-trunk. Rivulet2 has no hyper-parameter other than a foreground threshold, which can be trivially eliminated by the automatic thresholding methods in well-contrasted images. In our experiments, we found that the proposed algorithm was able to generate reasonable results in most of the challenging images acquired from various species and neuron types. Rivulet2 was shown to outperform Rivulet1 and several state-of-the-art methods in a majority of the bench-test images obtained from the DIADEM challenge and the BigNeuron challenge.

## II. METHODS

### A. Overview of Rivulet2

Taking a 3D grey scale image *I*(*x*) as the input with 3D coordinates *x*, a neuron tracing algorithm aims to output the neuron reconstruction as a tree graph model *G* where each tree node is assigned a 3D spatial coordinate and a radius. Each neuron tree node can have a degree between 1 and 3. The root node of *G* is defined as the soma centre.

To obtain this tree graph model, Rivulet2 starts with Gaussian and median filtering for very noisy images. A binary segmentation map *B*(*x*) is then generated to classify the voxels as foreground and background. The foreground voxels are considered as the potential neuronal signals imaged by light microscope. In practice, Rivulet2 is capable of generating reasonable results only with a coarse image segmentation generated by only applying a background threshold. We then generate a 3D boundary distance transform *DT*(*x*) based on *B*(*x*). The voxels close to the background have lower values than the voxels close to the neuronal centrelines in *DT*(*x*). Next, a time crossing map *T*(*x*) is generated with a fast marching method [27] using a speed image generated by *DT*(*x*). Based on the gradient of this map 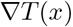, Rivulet2 traces each branch of the neuron tree iteratively. It starts with the geodesic furthest point remaining in the foreground and attempts to track back to the soma centre. An online confidence score is computed at each tracing step along with several other stopping criteria. The tracing iteration is stopped if any of the criteria is triggered. The area covered by the newly traced branch is marked on the time map *T*(*x*) to indicate that it has been explored. The newly traced branch is merged to the trunk if it touches the area covered by a previous branch. The whole process stops after all of the foreground areas have been explored. Finally, the short leaves and the unconnected branches are removed in the output tree *G* to ensure the neuron topology is valid for morphometric analysis.

### B. Time Crossing Map

The segmentation map *B*(*x*) is firstly obtained with a background threshold. We then use the multi-stencils fast marching (MSFM) [27] to obtain the geodesic distance between the soma centre *x*_*soma*_ and every voxel in the input image, including the background area. The fast marching method outputs a map of travelling time *T*(*x*) departs from the source point, *p*_*soma*_ in our case, to any voxel by solving the Eikonal equation

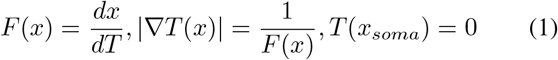

where *F* is the travelling speed defined at 3D coordinates *x*. To make the speed image *F*, we obtain a boundary distance transform *DT*(*x*). Each voxel of *DT*(*x*) contains its euclidean distance to the segmented boundary [3]. *argmax*_*x*_*DT*(*x*) and *maxDT*(*x*) are used as the soma centre *x*_*soma*_ and the soma radius *R*_*soma*_ respectively. Our speed image *F*(*x*) used in MSFM is formed as

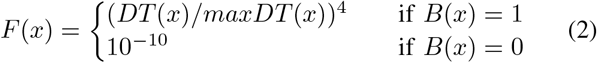

Thus, only the speed of the foreground area is determined by *DT*(*x*). The normalised *DT*(*x*) is powered by 4 to further highlight the centreline. We leave a small speed value 10^−10^ in the background area to allow the tracing to proceed when a gap presents. The background travelling speed would not outweigh the foreground speed, due to the large speed differences. MSFM is then performed on *DT*(*x*) with *x*_*soma*_ as the single source point. The computation of MSFM is stopped when all the foreground voxels with *B*(*x*) = 1 have been visited. Since the travelling time changes faster within the neuronal arbours than the background area, the gradient direction in 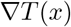 at each foreground voxel is expected to align with the orientation of the neuron arbour it resides in.

### C. Sub-Voxel Back-Tracking in a Single Branch

With the gradient descent on 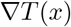, we can trace the neuron structure that a source node *p* resides in by repeatedly updating the location of *p* as

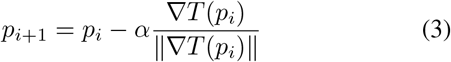

where *α* is the step-size constant. *p* is supposed to move from the outer area of the neuron towards the soma centre *x*_*soma*_. However, since most of the light microscopic images are under-sampled, the precision of voxel-wise gradient descent may introduce direction errors that affect the future tracing steps. Therefore, we use the sub-voxel gradient interpolation to perform the back-tracking with the fourth order Runge-Kutta method (RK4) as

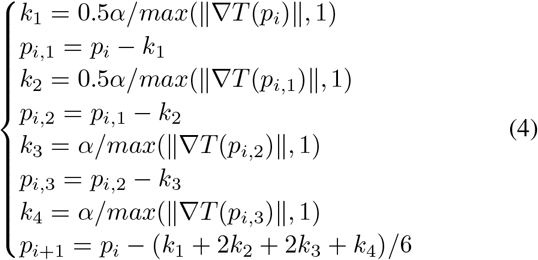

where *k*_1_, *k*_2_, *k*_3_, *k*_4_ are the direction vectors interpolated at the sub-voxel resolution. *α* is fixed as 1. To prevent tracing from stopping at a local minimal, the momentum is used instead for point update when the velocity 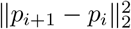 is small

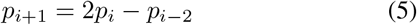

### D. Iterative Back-Tracking with Precise Branch Erasing

All the branches of a neuronal tree are traced iteratively, with the gradient back-tracking described in Section II-C. Next, we make a copy of *T*(*p*) that is denoted as *T*\*(*p*) for finding the starting point for each tracing iteration and labelling the traced branch. The values of the original *T*(*p*) are used for the branch erasing described later in this section. Each tracing iteration starts with the voxel *x*_*source*_ = *argmaxT*\*(*x*). *x*_*source*_ is considered as the location of either an undiscovered neuronal terminus or a noise voxel segmented by mistake. The position of the neuronal node *p* is updated by tracking from *x*_*source*_ to *x*_*soma*_ along the neuronal fibre curve *c*(*t*) that *x*_*source*_ might reside in using the RK4 tracking described in Eq.4. *c*(0) represents the start of the curve at *x*_*source*_ and *c*(1) represents the newly traced end of the curve. We track the distance *G*(*i*) that the *i*-th node has been travelled on the background as

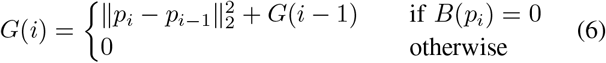

The radius *R*_*i*_ of the node at *p*_*i*_ is obtained by growing a spherical region centred at *P*_*i*_ as 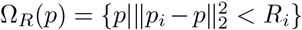 until 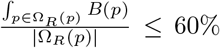, where |Ω_*R*_| is the volume of Ω_*R*_(*p*). Since the RK4 tracking is powerful of tracing across large gaps between the broken neuron segments, we designed a few stopping criteria to avoid Rivulet2 from generating false positives. The tracing of *c*(*t*) is stopped when any of the following criteria is triggered:

1. *c*(1) reaches the soma area when 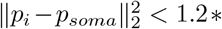 *R*_*soma*_
2. The online confidence (OC) score *P*(*c*(*t*), *B*(*x*)) is smaller than 0.2 or a deep OC valley is detected as described in Section II-E.
3. An exceptionally large gap presents in *c*(*t*) as described in Section II-E.
4. *c*(*t*) is ready to merge with another previously traced branch as described in Section II-F.
5. The tracing of *c*(*t*) has not moved out of the same voxel it reached 15 steps before.
6. *c*(*t*) reaches an out of bound coordinate.

To avoid repeatedly tracing the area covered by *c*(*t*) in the future iterations, *T*\*(*p*) is then erased as

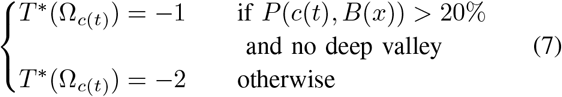

The erased regions Ω_*c*_(*t*) with *T*\*(Ω_*c*(*t*)_) = −1 is considered as erased by a neuronal fibre; it is otherwise considered as erased by a curve traced on the noise points. The regions with *T*\*(*x*) < 0. are thus excluded for selecting new *x*_*source*_ in future iterations. The erased regions also indicate when a newly traced branch should be merged as described in Section II-F. At the end of a tracing iteration, *x*_*source*_ = *argmax*_*x*_*T*\*(*x*) is chosen from the remaining *T*\*(*x*) as the location of the new source point *p* for the next iteration. The entire algorithm terminates when all the foreground region has been erased from *T*\*(*x*).

The estimate of Ω_*c*(*t*)_ is important for tracing accuracy as well as the running time. Rivulet1 [8], [26] used a similar method for region estimation as the pruning based methods [4], [28] by forming it as the union region Ω_*R*_ of all the spherical regions covered by the nodes in *c*(*t*)

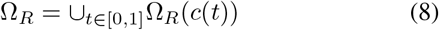

However, since Ω_*R*_was only an approximated estimate, when Ω_*R*_(*c*(*t*)) is locally over-reconstructed, there is a risk that voxels on the unexplored branches and the branch forking might be erased; Otherwise, it leaves small fragments remaining at *T*\*(*p*), resulting in more tracing iterations and over-reconstructed branches.

In Rivulet2, we form a new region Ω by combining another region generated with the original time map *T*(*p*)

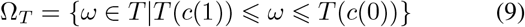

A region Ω_*R\**_ that is slightly larger than the previous Ω_*R*_ is formed with 120% × *R*_*i*_ to include all the possible candidate voxels to be erased. Ω is then formed as Ω = Ω_*R\**_ ⋂ Ω_*T*_. The formulation of Ω
 is illustrated in Fig.2(d). Ω is a precise estimate of the covered region of *c*(*t*) by considering the travelling time generated by MSFM. In a majority of cases, Ω covers exactly the branch boundary without leaking at the branch forking points.

**Fig. 2.**
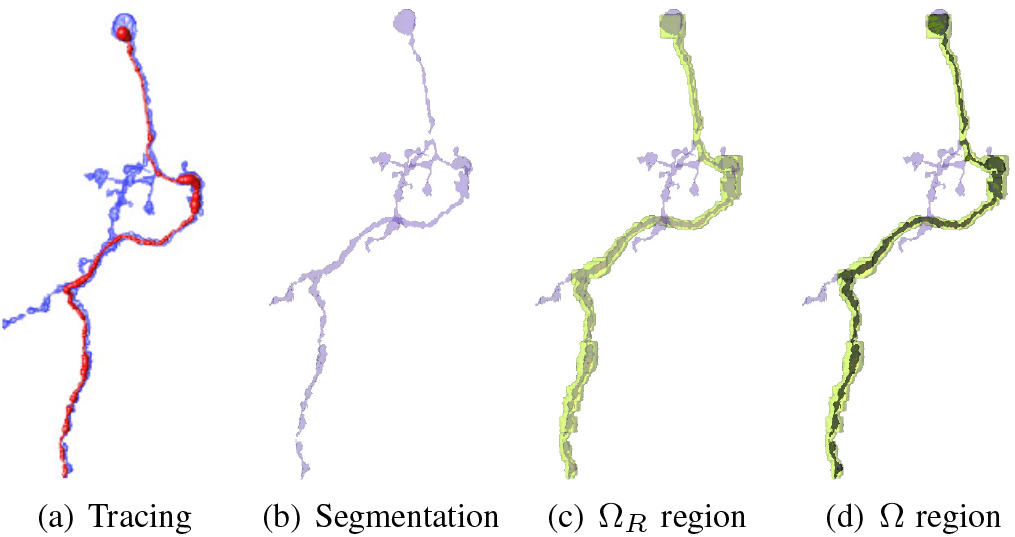
Illustration of the contour used for branch erasing. (a) is the tracing of one iteration (red) overlaid on the original image (blue); (b) is the segmentation used for Rivulet2 tracing; The green area in (c) is the Ω_*R*_ region which is also used to erase the traced branch in Rivulet1; The black area inside Ω_*R*_ in (d) is the region Ω used in Rivulet2. Since Ω enables a more accurate estimate of the traced region, Rivulet2 traces the entire neuron faster than Rivulet1 without breaking the connection at the neuronal joints.

### E. Branch Cut with Online Confidence Score

Since *x*_*source*_ can sometimes be a noise voxel, an effective method is needed to distinguish branches traced on neuronal fibres and the ones traced from noise voxels. Rivulet1 uses a single gap threshold to stop the tracing when a certain number of steps have been made in the background. However, the choice of the gap threshold is ill-posed. For Rivulet2, we compute an online confidence (OC) score *P*(*c*(*t*), *B*(*x*)) for each tracing step. OC is defined as the proportion of backtracking steps that are made in the foreground voxels so far

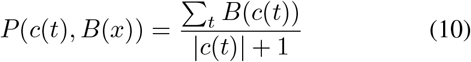

where ∑_*t*_*B*(*c*(*t*)) represents the number of steps in the foreground; |*c*(*t*)| is the number of total steps in *c*(*t*). Different OC curves generated in a single fly neuron are shown in Fig. 3(a). *P*(*c*(*t*), *B*(*x*)) is expected to decrease quickly during back-tracking if the tracing starts from a noise point. *P*(*c*(*t*)) would otherwise remain a high value if the back-tracking jumps over small neuron fibre gaps since the majority of back-tracking steps are made in the foreground voxels. The +1 term ensures *P*(*c*(*t*), *B*(*x*)) on a noise branch starts from 0:5 at its first step. The back-tracking is stopped if *P*(*c*(*t*), *B*(*x*) is lower than 20% as shown with the horizontal line in Fig. 3(a), indicating it was tracked from a noise voxel far away from the neuron fibre. The regions erased by low confidence branches are considered to be noise regions. The low confidence branches are excluded from the final neuronal tree. It is also notable that the future iterations are allowed to trace across the regions with *T*\*(*p*) = −2 without the branch merging being triggered. The branches with *P*(*c*(*t*)) > 20% might show a dramatic decrease at the beginning and an increase after it reaches the neuron fibre if it is traced from a noise voxel. As depicted in Fig. 3(a), deep valleys would appear along the OC curves of the noisy branches, indicating the step when it touches the neuron fibre. We erase *T*\*(*p*) with only the former part of the branch with −2 before the valley if *P*(*c*(*t*), *B*(*x*)) < 50% at the valley point.

**Fig. 3.**
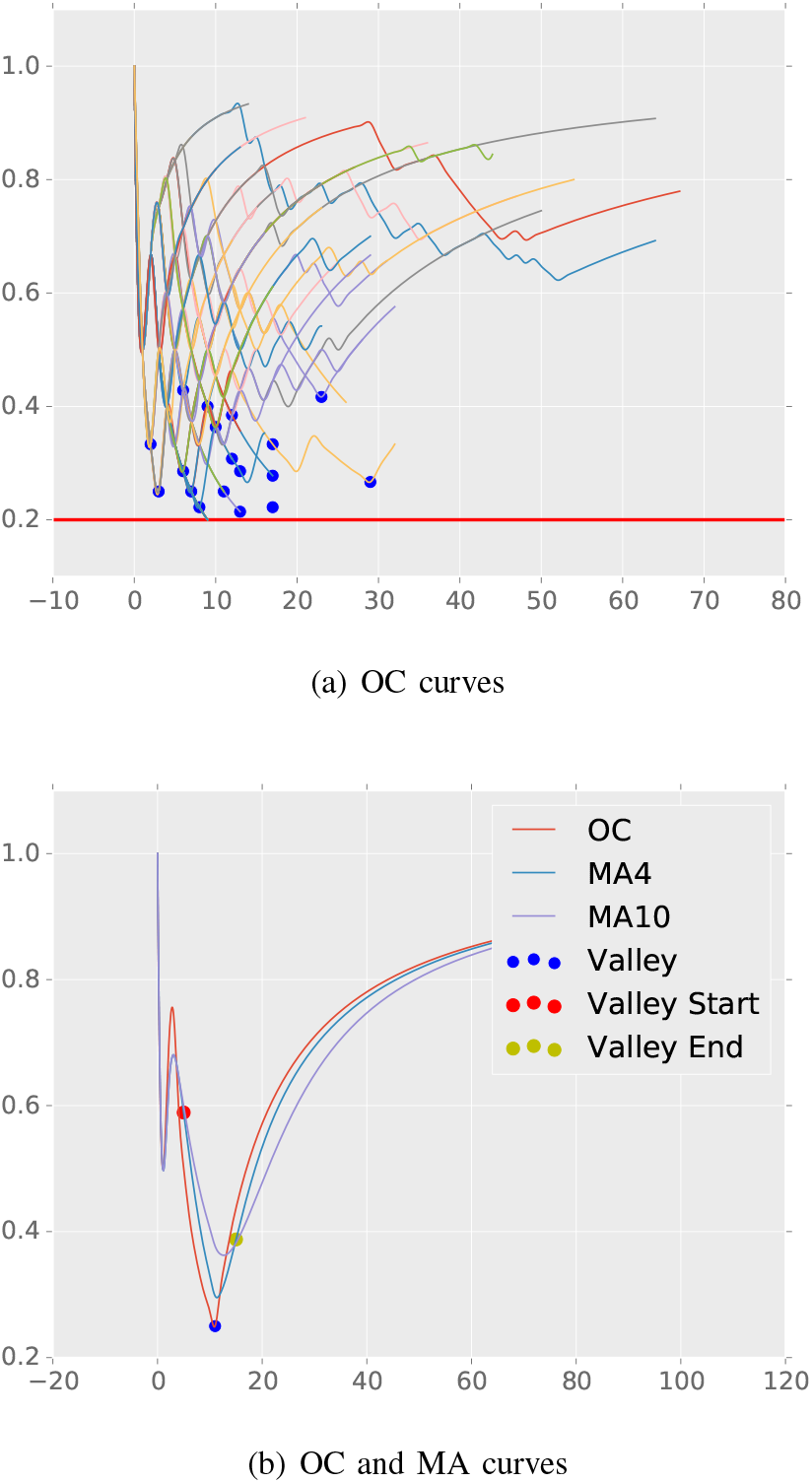
(a) visualises the online confidence (OC) curves while tracing a single neuron cell from a noisy image. Some of the tracing iterations are stopped when their OC curves touch 0:2 (the red horizontal line). For the tracing iterations with OC scores higher than 0:2, the branches traced before the deep valleys, represented by blue spots, are discarded. (b) shows a single OC curve accompanied by two of its moving average (MA) curves with the window sizes 4 and 10. Inspired by a financial analysis technique, the deep valley of OC curve is detected at the lowest value between the two crossings of the MA curves.

When the image is highly noisy, it might be insufficient to identify a deep OC valley with only the lowest value of *P*(*c*(*t*), *B*(*x*)) across the entire branch. We use the exponential moving average (EMA) that is widely used in the financial analysis to detect the deep OC valleys. The EMA is defined as

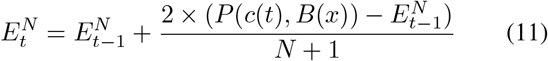

where *E*_*t*_ is the EMA score with the window size of *N* at the step *t*. We use two different window sizes 4 and 10 to track a short-term EMA 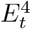 and a long-term EMA 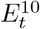. The valley point is found at the lowest point in *P*(*c*(*t*), *B*(*x*)) between the two crossings of 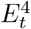 and 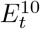 if such two crossings exist. The example valley points are shown as the blue spots in Fig. 3(a).

Using both the bottom boundary and the valley points, the OC score is a simple but effective approach to identify most of the neuronal gaps and the noise points. However, some of the images also contain bright curvilinear structures that do not belong to the same neuron cell of interest. For example, the single neurons extracted from the Brainbow [29] images with colour extraction sometimes contain fibres of other neurons as shown in Fig 1(e). Though the gaps between such fibres and the neuron of interest are normally large, *P*(*c*(*t*), *B*(*x*)) could remain high. To stop tracing from such irrelevant fibres, the tracing stops when a continuous gap *G*(*t*) is larger than 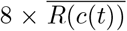 where 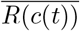 is the mean radius estimated on *c*(*t*)

### F. Branch Merging

When the branch *c*(*t*) reaches a voxel *x* with *T*\*(*x*) = −1, it means the branch has reached an area explored by the previous iterations. Rivulet1 stops the tracing iteration immediately in such voxel and search for a previously traced node to connect. However, it may cause topological errors since the endpoint of *c*(*t*) might still be far from the branch that it should be merged into. In Rivulet2, the tracing iteration does not stop once it touches the boundary of a previously traced area. Instead, it keeps performing back-tracking after the boundary touch and seeks for a candidate node from the previous branches to merge at each step. It is merged into the tree trunk if the closest node *p*_*min*_ is either 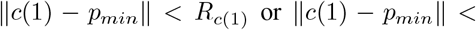 *R*_*pmin*_. The wiring threshold that controls the tolerable node distance to merge used in Rivulet1 is thus no longer needed in Rivulet2.

### G. Post-processing

After all the back-tracking iterations, only the largest connected section is kept. The majority of the discarded branches are the background bright curvilinear structures that do not belong to the same neuronal cell. It is also optional to remove short leaves having the lengths shorter than 4 as long as spine detection is not required. Though the detection of the node type is normally not required in the challenges such as DIADEM [1] and BigNeuron [2], the node types such as soma, fork points, end points are labelled when the branch is added to the tree trunk. It is not capable of distinguishing the fibre classes including apical dendrites, basal dendrites and axons.

## III. EXPERIMENTAL RESULTS

### A. Data

The images used in this study were recruited from both the DIADEM challenge [1] and the BigNeuron challenge [2]. Nine image stacks of Olfactory projection (OP) fibres were obtained from the DIADEM challenge. The OP dataset was widely used and compared in the previous studies. Each OP image stack contains a separate drosophila olfactory axonal projection with a corresponding gold standard reconstruction. The OP images were acquired with 2-channel confocal microscopy. The groundtruth manual tracings were obtained with Neurolucida (Williston, VT) and Amira (Chelmsford, MA) extension module hxskeletonize [30]. We manually fixed some of the incomplete manual reconstructions with Vaa3D before benchmarking all the methods.

We use images from the BigNeuron project as the second dataset with a larger diversity of neuron types and image conditions. The BigNeuron images were recruited from different neurobiology labs globally. Each image was traced and validated by at least three neuroscientists using Vaa3D [31]. We chose nine challenging subsets of the BigNeuron cohort with 114 images in total, containing neuron cells from different species including fly, fruit fly, human, zebrafish, silkmoth, frog and mouse. The subsets were chosen considering the feasibility for automatic large-scale bench-marking. Also, to evaluate the robustness of Rivulet2 on large-scale image datasets, we tested it against the first-2000 dataset containing 2000 fruit fly neuron images.

### B. Implementation and Evaluation

To quantitatively evaluate Rivulet2, we used the Python implementation Rivuletpy ^1^ released together with this paper. A C++ implementation of Rivulet2 is also available in Vaa3D as a neuron tracing plugin ^2^. We compared Rivulet2 with several state-of-the-art neuron tracing methods, including APP2 [28], SmartTracing (SMART) [5], Farsight Snake (SNAKE) [16], Probability Hypothesis Density Filtering (PHD) [11], Ensemble Neuron Tracer [10], Neutube [7] and its predecessor Rivulet1 ([8], [26]). We used the Vaa3D ported implementations for bench-marking the methods APP2, SMART, SNAKE, ENT, and Neutube. For APP2, we used gray-weighted distance transform (GWDT) and disabled the automatic image resampling. For PHD, we used its FIJI implementation and performed grid-search for hyper-parameters using the FIJI batch-processing macro provided together with the plugin. The Rivulet Matlab Toolbox^3^ was used for testing the performance of Rivulet1. We use grid search for the wiring and the gap thresholds for Rivulet1. The same manually selected background thresholds were used for evaluating all the compared methods when required.

NeuroM (https://github.com/BlueBrain/NeuroM) is used to validate the outputs before obtaining the quantitative analysis. The empty or invalid neurons were not included in the quantitative results. We use the precision, recall and F1-score to evaluate the geometric appearance of the automated reconstructions. A node in the automatic reconstruction is considered as a true positive (*TP*) if a ground truth node can be found within four voxels; it is otherwise a false positive (*FP*). A false negative (FN) is defined when there is no automatically reconstructed node within four voxels of a ground truth node. The precision is defined as *TP*/(*TP* + *FP*), and the recall is defined as *TP*/(*TP* + *FN*). The F1 score balances the precision and recall as 2 × *precision* × *recall*/(*precision* + *recall*). Also, we compute the node distance measurements proposed in [32] which are the spatial distance (SD), significant spatial distance (SSD) and the percentile of distant spatial nodes (SSD%). SD measures the mean distance between each pair of closest nodes between two neuron reconstructions. SSD measures the SD distance between each pair of closest nodes when they are at least two voxels away from each other; SSD% measures the percentile of the reconstructed nodes that are at least two voxels away. All the bench-marking were performed using the Artemis high-performance computing (HPC) infrastructure at the University of Sydney.

### C. Diadem Results

We show the Rivulet2 reconstructions of all 8 compared neurons in the DIADEM OP dataset in Fig. 4. The manual reconstructions are shown on the left, and the Rivulet2 (R2) reconstructions are shown on the right. The proposed Rivulet2 algorithm can obtain visually identical reconstructions to the groundtruth tracings by using a fixed threshold of 30 on all images. The quantitative results are shown in Fig. 5 with box plots. Rivulet2 outperforms the compared methods in precision, F1 score, SD, SSD and SSD%. A small drop of recall is seen compared to Rivulet1 since Rivulet1 tends to generate more false positive branches when noise presents. The robustness of Rivulet2 can also be seen in the relatively small variances in each metric.

**Fig. 4.**
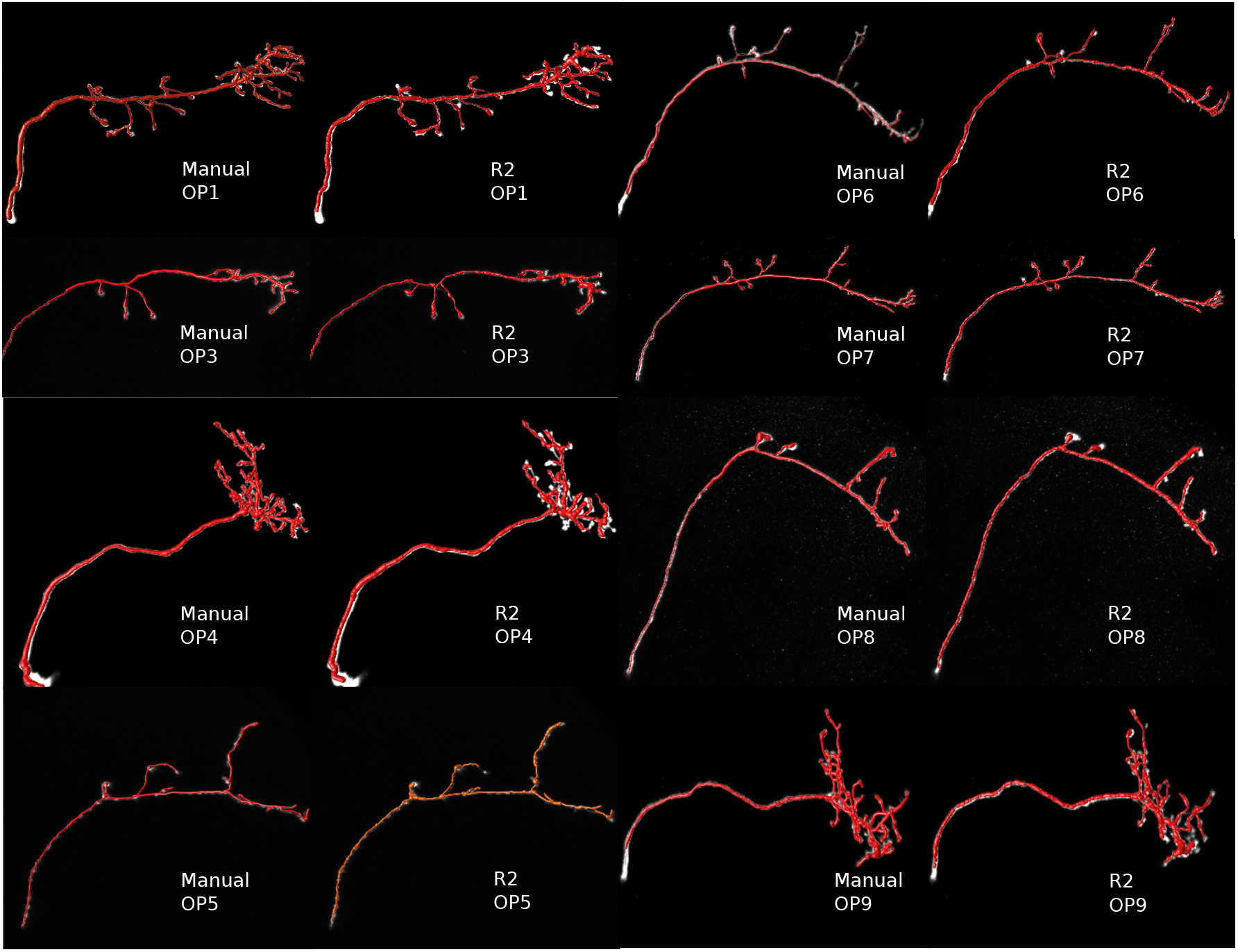
The visual inspections of the eight neurons in the OP neurons. The manual reconstructions are shown on the left, and the Rivulet2 (R2) reconstructions are shown on the right. The reconstructions are overlaid on the original images. The proposed Rivulet2 algorithm can obtain visually identical reconstructions to the groundtruth tracings.

**Fig. 5.**
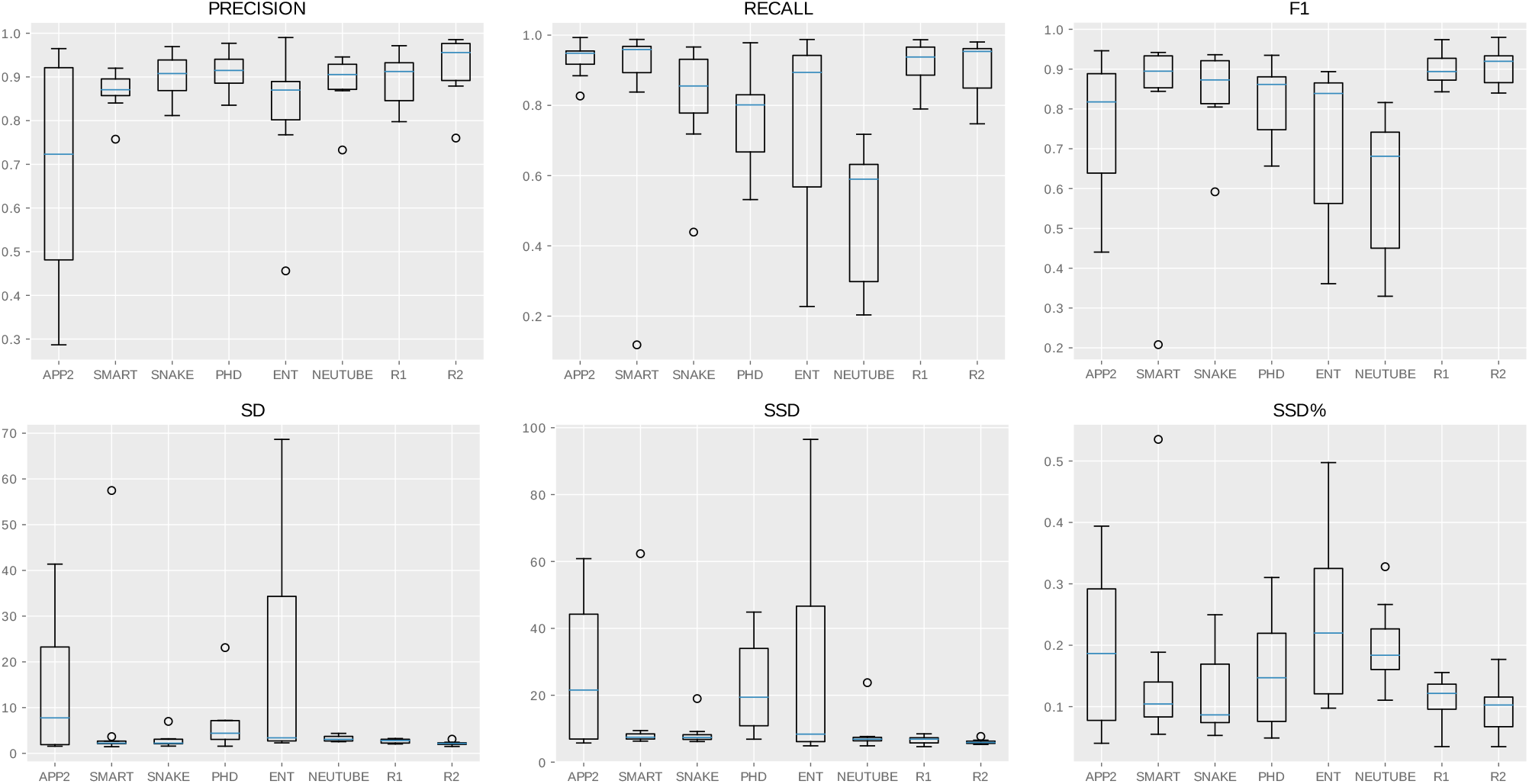
The quantitative results of the OP dataset containing 8 image stacks.

### D. BigNeuron Results

We selected two challenging images to visually compare the results shown in Fig. 6 and Fig. 7. The neuron in Fig. 6 is a fly neuron with dense noise in the background. Both Rivulet1 and Rivulet2 were able to reconstruct the entire neuron without being interrupted by the noise. Comparing to Rivulet1, Rivulet2 was able to suppress the majority of the false positive branches. Fig. 7 shows a zebrafish adult neuron with many gaps in the background containing strong noise. There are also many irrelevant curvilinear structures residing in the background that are hard to eliminate when only local information is considered. Rivulet2 could reconstruct reasonable results across the entire neuron with little false positive branches. Rivulet1 generated many redundant segments due to the noise and the irrelevant bright area on the top-left corner.

**Fig. 6.**
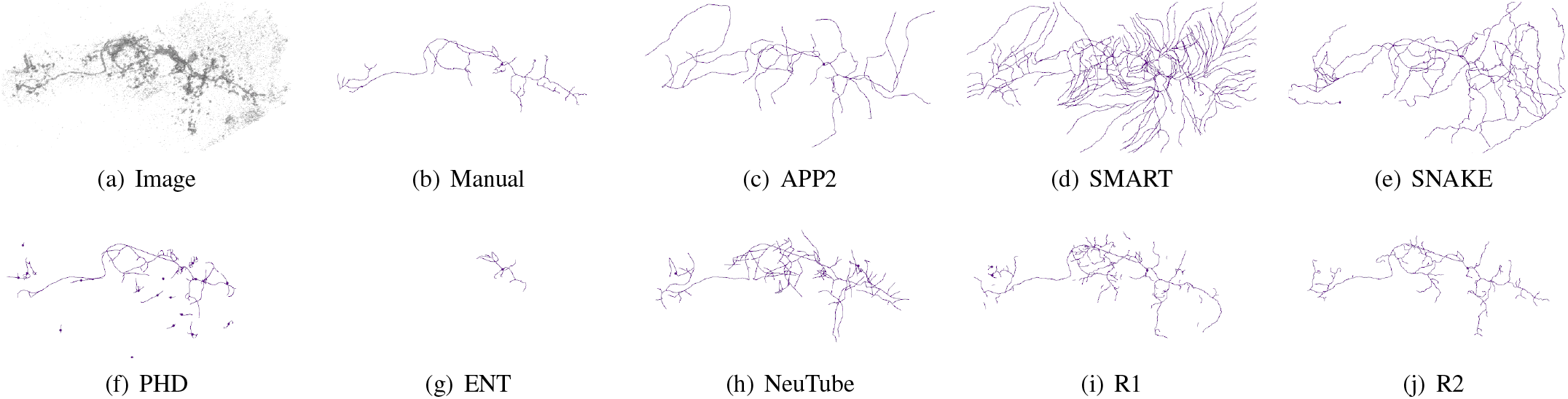
The visual inspection of a fly neuron from the BigNeuron project. The image is rendered with inverse intensities to make the image noise visible in low-resolution. The reconstructions are visualized with Vaa3D using the line-mode.

**Fig. 7.**
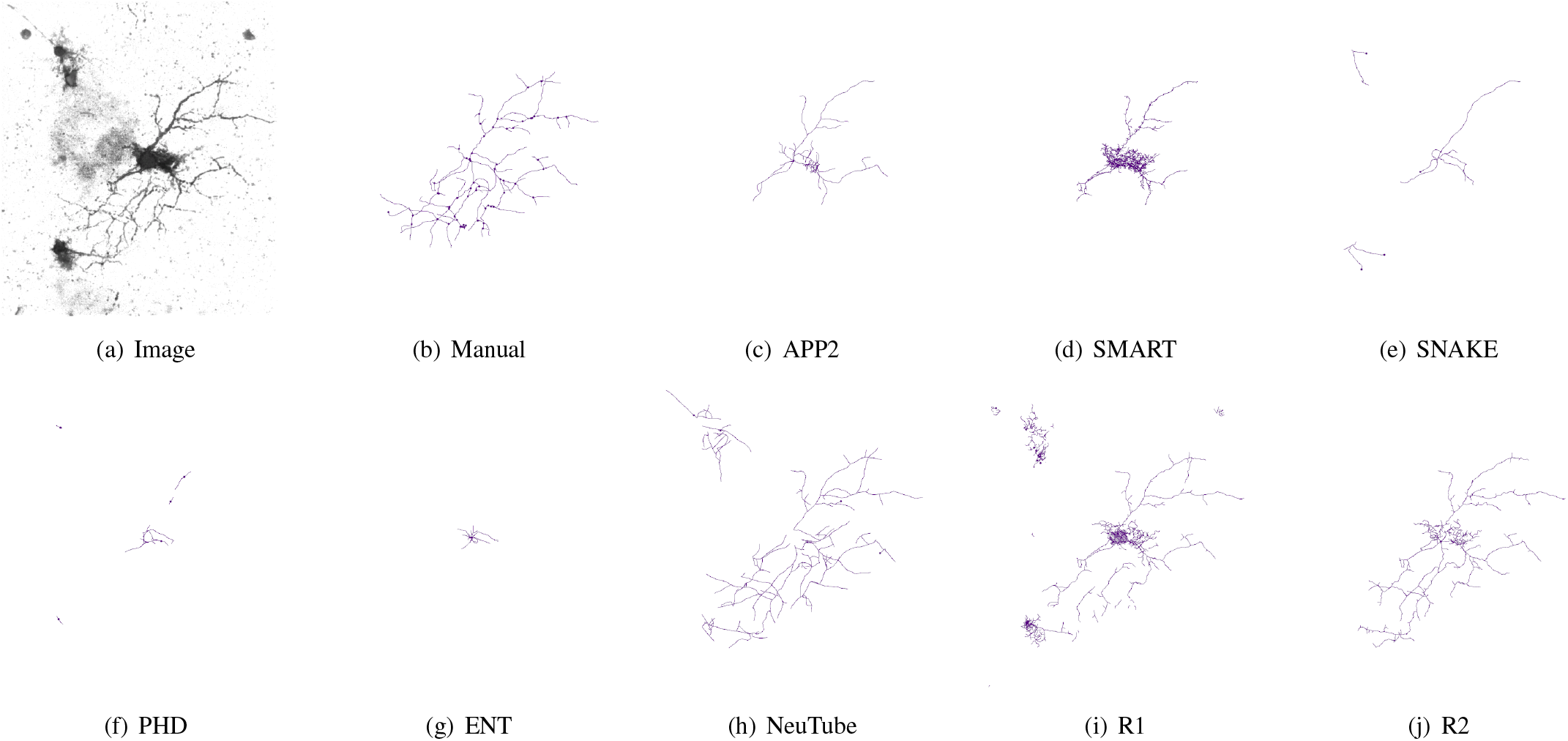
The visual inspection of an adult zebrafish neuron from the BigNeuron project. The image is rendered with inverse intensities to make the image noise visible in low-resolution. The reconstructions are visualized with Vaa3D using the line-mode.

We quantitatively compared all the methods with the gold standard manual reconstructions traced by the BigNeuron community as shown in Fig. III-B. Similar to the result in the OP dataset, Rivulet2 achieved the highest precision, F1-score and slightly lower recall than the Rivulet1 and SmartTracing. At the same time, Rivulet2 obtained the lowest or comparable values in the distance metrics (SD, SSD and SSD%).

To test the robustness of the proposed method on batch-processing of large-scaled datasets, we applied it to the first-2000 dataset released by the BigNeuron project that contains 2000 neurons which have been coarsely segmented. Since this dataset was not manually annotated, we use it only for comparing the running time of Rivulet1 and Rivulet2. We did not benchmark the running time of the other state-of-the-art methods since they were implemented in different languages. The neurons with the top eight total dendrite lengths are shown in Fig. 9. The resulted nodes were sorted by the Vaa3D Sort SWC plugin and validated by NeuroM. 1997 out of 2000 reconstructions were validated neuron trees. We manually inspected the three failed neurons and found the failures were only caused by 3 broken images in this repository. The average running time of Rivulet2 is 110.875 seconds which is more than four times faster than Rivulet1 (456.605 seconds). The speed increase is mainly introduced by the precise branch erasing and the online confidence score. Rivulet2 is slower than the C++ implementation of APP2 (14.950 seconds) mainly due to the gradient interpolations needed in the sub-voxel backtracking, and the MSFM performed across the entire image.

**Fig. 8.**
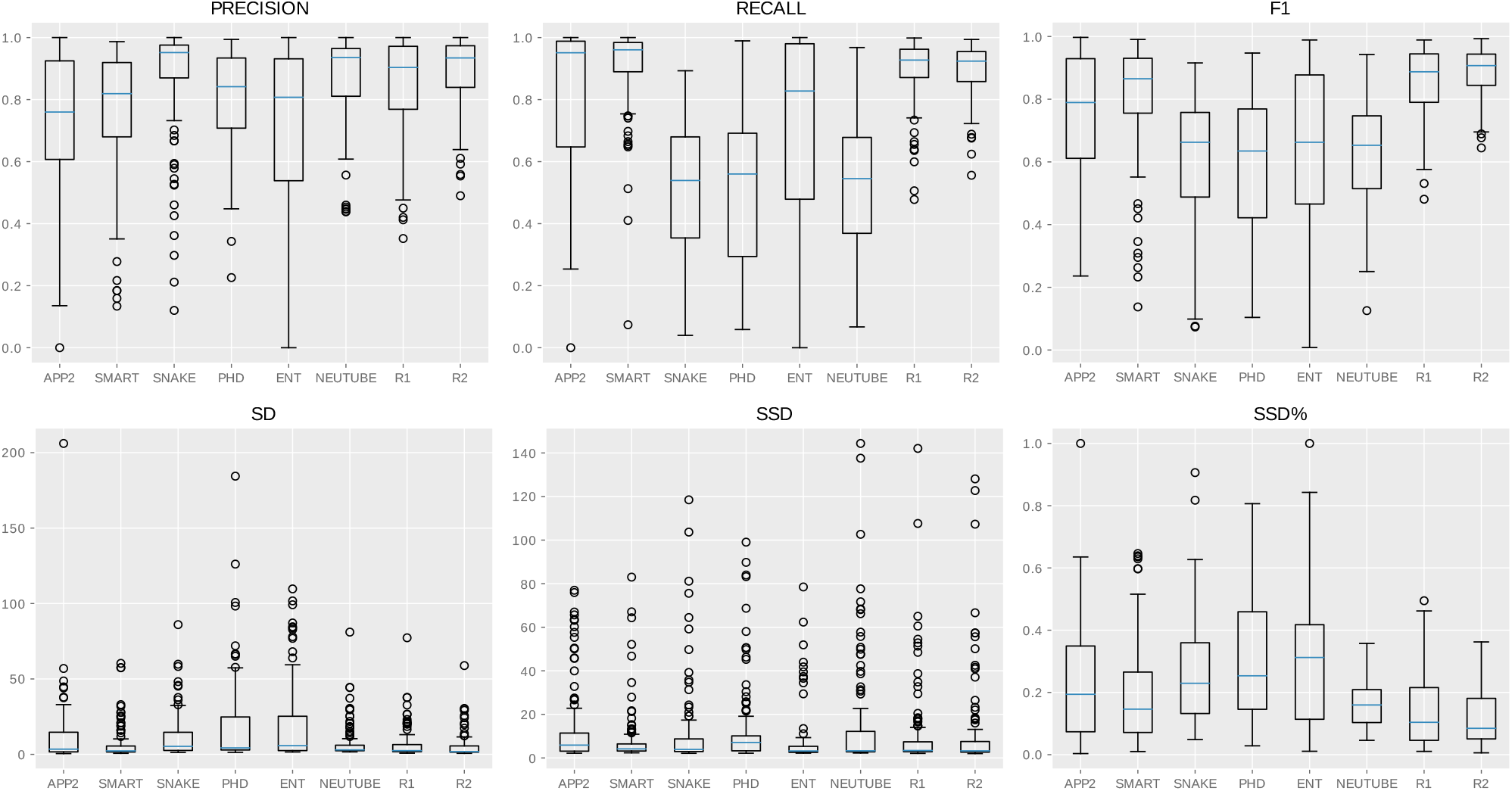
The quantitative results of the BigNeuron dataset containing 114 image stacks.

**Fig. 9.**
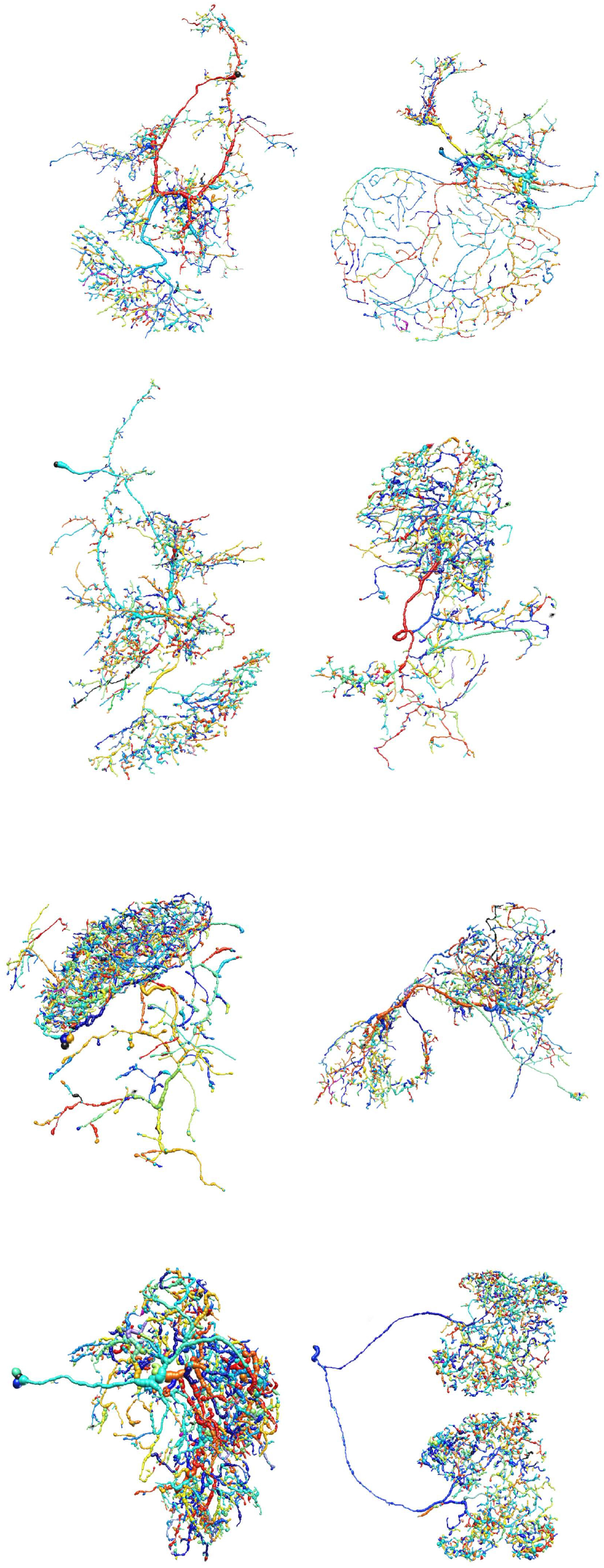
The Rivulet2 reconstructions of the top 8 neurons in the first-2000 dataset regarding the number of nodes. The first-2000 dataset was released by the BigNeuron project.

## IV. CONCLUSION

In this study, we proposed a fully automatic 3D neuron reconstruction method Rivulet2. By evaluating the proposed method with the newly released data from the BigNeuron project, we showed that Rivulet2 was capable of generating accurate neuron tracing results in most challenging cases with only a single background threshold. Rivulet2 was also capable of producing topologically authentic neuron models for morphometrics analysis. Comparing to Rivulet1, it is approximately four times faster. It also outperformed state-of-the-art neuron tracing algorithms on most of the selected BigNeuron benchmark datasets.

## ACKNOWLEDGEMENT

We thank the BigNeuron community for providing the data and the discussions. The authors acknowledge the Sydney Informatics Hub for providing the Artemis computing clusters that were used in the evaluation. Siqi Liu is supported by the Australian Postgraduate Award (APA) scholarship and the Google PhD Fellowship in computational neuroscience. This work is also supported by the Allen Institute for Brain Science.

